# *Staphylococcus aureus* Persistence in Osteocytes: Weathering the Storm of Antibiotics and Autophagy/Xenophagy

**DOI:** 10.1101/2023.10.27.564475

**Authors:** Nicholas J. Gunn, Anja R. Zelmer, Stephen P. Kidd, Lucian B. Solomon, Dongqing Yang, Eugene Roscioli, Gerald J. Atkins

**Author notes:** These authors share senior author status (Eugene Roscioli, Gerald J. Atkins).

## Abstract

**Background:** *Staphylococcus aureus* is a major causative pathogen of osteomyelitis. The intracellular infection of osteocytes and related bone cells can persist despite application of gold-standard clinical interventions. The mechanisms by which intracellular *S. aureus* persists during antibiotic therapy are unknown. In this study, we apply *S. aureus* to an *in vitro* model of differentiated osteocytes to investigate whether antibiotic-mediated dysregulation of autophagy contributes to this phenomenon.

**Methods:** Human osteocyte-like cells were exposed to combinations of rifampicin, vancomycin and modulators of autophagy, in the presence or absence of *S. aureus*. Intracellular bacterial growth characteristics were assessed through CFU analysis, viable bacterial DNA abundance and the rate of escape into antibiotic-free medium, in parallel with measures of host cell autophagic flux.

**Results:** Rifampicin, alone or in combination with vancomycin, caused a rapid decrease in the culturability of the intracellular bacterial community, concomitant with stable or increased absolute bacterial DNA levels. Both antibiotics significantly inhibited autophagic flux. However, while the modulation of autophagic flux affected bacterial culturability, this modulation did not affect viable bacterial DNA levels.

**Conclusions:** Autophagy was shown to be a factor in the host-pathogen relationship in this model, as its modulation affected the growth state of intracellular *S. aureus* with respect to both their culturability and propensity to escape the intracellular niche. Whilst rifampicin and vancomycin treatments moderately suppressed autophagic flux acutely, this did not explain the paradoxical response of antibiotic treatment in decreasing *S. aureus* culturability while failing to clear bacterial DNA and hence intracellular bacterial load. Thus, whilst rifampicin and vancomycin exhibited off-target effects that modulated autophagy in osteocyte-like cells, this could not explain the persistent infection observed for *S. aureus*.

## INTRODUCTION

*Staphylococcus (S.) aureus* is the most common causative pathogen in all forms of adult osteomyelitis, exemplified by that associated with periprosthetic joint infection (PJI). These infections can become refractory to antibiotic therapies through defined resistance mechanisms, or exhibit antibiotic tolerance (in the absence of resistance), by adopting alternative growth phenotypes (1, 2). It has been demonstrated that intra-osteocytic infection of *S. aureus* is a feature of osteomyelitis, including in PJI, and that *S. aureus* adapts to its intracellular environment by forming small colony variants (SCV) (3–6). Such phenotypic adaptation is consistent with the high incidence of post-treatment reinfection in PJI, despite antibiotic therapy, as high as 8.8-35.5% within two years using standard approaches (7–11).

Autophagy is a fundamental cellular process with myriad functions related to immunity and cellular homeostasis (12, 13). Crucially, autophagy is central to the process of degrading and recycling damaged or unnecessary cellular components. LC3 (Microtubule-associated protein 1A/1B-light chain 3) is pivotal in the function of autophagosomes that engulf cellular material targeted for degradation. Initially present in the cytosol as LC3-I, it undergoes lipidation to become LC3-II, which is then recruited to the growing autophagosomal membrane. This recruitment is essential for autophagosome elongation and closure, and facilitates cargo sequestration within the autophagosome. Sequestosome 1 (SQSTM1), also known as p62, acts as a selective autophagy receptor, recognising ubiquitinated cargo destined for degradation and linking it to LC3-II on the autophagosomal membrane. Xenophagy is a specialised form of autophagy that specifically eliminates invading pathogens, such as bacteria, viruses and parasites. Degradation of the autophagosomal cargo is mediated by fusion with lysosomes to form the autophagolysosome (14–17).

There are few situations, in which autophagy/xenophagy is subject to clinical or diagnostic inquiry, despite the critical role played by this pathway in host cell clearance of intracellular infections. Given that pathogenic bacteria have evolved means to subvert, disrupt or even make use of components of the autophagic/xenophagic pathway to reside intracellularly, it is important that antibiotics do not contribute to this outcome (18). We have shown that antibiotics can potently arrest innate intracellular bacterial clearance processes in lung epithelium (19). Several antibiotics used for the treatment of Staphylococcal PJI (20) have been identified to elicit this off-target effect (5). For example, the aminoglycoside gentamicin, which is commonly used in antimicrobial-loaded bone cements, has been demonstrated to inhibit autophagosome-lysosome fusion, an effect that is also mediated by intracellular *S. aureus* (21, 22). Other aminoglycosides, including tobramycin and daptomycin, when fully protonated within the acidic environment of lysosomes aggregate and inhibit of lysosomal phospholipase activity (23). In addition, vancomycin-induced kidney injury appears to be at least partly mediated by autophagic dysregulation, via its accumulation and subsequent rupture of autophagolysosomes (24). Vancomycin has also been demonstrated to block autophagic flux in macrophage cell lines (25). Rifampicin, on the other hand, has been shown to inhibit rapamycin-induced autophagy through the suppression of the positive regulator of autophagy, PP2A (26).

Taken together, these observations point to a scenario whereby the efficacy of antibiotics in clearing extracellular infection may mask undesirable off-target effects on autophagy/xenophagy that promote intracellular residence of *S. aureus*. In the current study, we sought to determine the involvement of autophagy in a human osteocyte model of *S. aureus* infection, and the impact of two commonly co-administered antibiotics, rifampicin and vancomycin, on this interaction. We found significant interruption of normal autophagy by both antibiotics but this may not be consequential to the capacity of *S aureus* to persist within osteocytes.

## MATERIALS & METHODS

### Cell Culture

Cell culture of SaOS2 cells and differentiation under osteogenic growth conditions to an osteocyte-like stage (SaOS2-OY) was performed as previously reported (24, 25). Briefly, SaOS2 cells were cultured for 28 days in differentiation medium consisting of αMEM (Gibco, NY, USA) with 10% v/v foetal calf serum, 10 mM HEPES, 2mM L-Glutamine (Thermo-Fisher, VIC, Australia) and 50 µg/ml ascorbate-2-phosphate and 1.8 mM potassium di-hydrogen phosphate (Sigma, MO, USA). Media were refreshed twice weekly during the 28 day differentiation period, and weekly thereafter. During differentiation the medium was supplemented with penicillin/streptomycin (Thermo-Fisher, VIC, Australia), both at 1 unit/ml. Following infection, where relevant, media were either antibiotic-free or supplemented with 10µg/ml lysostaphin (AMBI Products LLC, NY, USA). Routine light microscopy showed that the differentiated osteocyte-like cultures exhibited the characteristic mineralised extracellular matrix consistent with this model (27, 28).

### Bacterial Culture and SaOS2-OY Infection

Growth and quantification of the methicillin-resistant *S. aureus* (MRSA) strain, WCH-SK2, was performed as described (27). This strain was chosen due to its demonstrated ability to establish intracellular infections in human osteocyte-like cells and in so doing rapidly alter their growth phenotype, in particular switching to a SCV state (6, 27). Briefly, bacteria were streaked onto nutrient broth agar (NBA) from glycerol stocks to ensure the culture was a monoculture of the intended strain. Isolated colonies were sub-cloned twice into nutrient broth. Log-phase broth was pelleted (2000 x g, 10 min, ambient temperature) and resuspended in phosphate buffered saline (PBS) to approximate the desired target CFU/ml from a standard curve of absorbance at OD_600nm_.

Infection of SaOS2-OY cultures was performed, as described (27). Briefly, differentiation medium was removed from SaOS2-OY cultures and the wells washed twice with PBS before the addition of the suspended bacteria. The actual CFU/ml was confirmed by plating serial dilutions of the inoculate onto NBA. Following a 2-hour infection period, the wells were washed twice with PBS and exposed to medium containing 10 µg/ml lysostaphin for 2 hours to clear extracellular bacteria. For ‘acute infection’ experiments, media were replaced with fresh media containing the described treatments. For ‘chronic infection’ experiments, cultures were maintained in media containing lysostaphin for a period of 8 days, whereupon cell lysates were non-culturable prior to commencement of treatments.

### Quantification of Bacterial Burden by qPCR

For intracellular bacterial retrieval, host cells were lysed in sterile water for 20 minutes at 37°C. Where exclusively viable bacterial DNA was required, the lysate was treated with the photoreactive DNA-binding dye PMAxx (Biotium, CA, USA). Briefly, PMAxx was added to lysates at a final concentration of 20 µM and this was incubated for 10 minutes at 37°C under blue light. After this, DNA was extracted using the Gram-positive bacterial cell lysate protocol as per the PureLink^TM^ genomic DNA mini kit (Thermo-Fisher), except replacing 20 µg/ml lysostaphin for the suggested lysozyme. Real-time qPCR was performed to quantify relative bacterial gene abundance with RT^2^ SYBR Green Fluor qPCR Mastermix chemistry (Qiagen, Limburg, Netherlands) and the CFX Connect thermocycler (BioRad, CA, USA). Where appropriate, bacterial DNA quantity was normalised to the magnitude of human *ACTB* DNA using the 2^-ΔCt^ method. When PMAxx treatment was used, as it inhibits extraction of human DNA from the lysates, human DNA abundance was determined from replicate cultures extracted in the absence of PMAxx. The sequences of the oligonucleotide primer sets used are listed in **Supplementary Table 1.**

### Quantification of bacterial gene expression

Bacterial transcripts were isolated using the Trizol reagent method (Life Technologies, NY, USA) and RNA reverse transcribed using the iScript gDNA clear cDNA synthesis system (Biorad, CA, USA). qPCR of complementary DNA was performed as previously described and normalised to human *ACTB* mRNA quantity using the 2^-ΔCt^ method (6).

### Measurement of SaOS2-OY cell viability

Cell viability was assessed using the lactate dehydrogenase (LDH) enzyme activity method (Cytotoxicity Detection Kit version 11; Sigma-Aldrich) to quantify compromised cells in response to exposure to antibiotics. SaOS2-OY cultures seeded into 48-well tissue culture plates were treated with a dose range of either rifampicin (0.75-75 µg/ml) or vancomycin (3-800 µg/ml) diluted in differentiation medium. LDH activity quantification was performed 24 hours after the initiation of treatments, following the manufacturer’s protocol.

### Measurement of Culturable Bacteria per Well

Bacterial outgrowth was routinely monitored by observation of media turbidity. For experiments where bacterial outgrowth was an outcome, this was further measured by daily plating of 1.5 µl of media onto NBA to enumerate viable cells. For the purposes of measuring culturable intracellular bacteria per well, after thorough rinsing in sterile media to remove antimicrobial agents, SaOS2-OY cells were lysed by 20 minute exposure to pure water at 37°C. Following lysis, the cell layer was scraped into solution and agitated by repeated passage through a 200 µl pipette tip. The resulting clumped, cell debris was separated by centrifugation from the lysate which was serially diluted 1/10 in sterile PBS and 100 μl plated onto NBA and incubated at 37°C. Viable cells were counted after 48 hours of incubation. All plates were left for a minimum of a further 14 days in the incubator to ensure that all culturable bacteria have time to generate macroscopically visible colonies.

### Antibiotic treatments

When analysing and comparing the effects of antibiotics on bacterial outcomes we used minimum bactericidal concentrations (MBC) for planktonic growth to normalise the comparison. These were determined by octuple replicate culture of log-phase bacteria into fresh nutrient broth containing a dose range of the antibiotic in question. The lowest dose for which no growth occurred following 24 h culture was designated as the MBC. Culture was confirmed by drop plating from the broth onto nutrient broth agar. For vancomycin, the planktonic MBC for *S. aureus* WCH-SK2 was measured to be 8 µg/ml. The equivalent concentration for rifampicin was 0.75 µg/ml. When interrogating the effects of antibiotics on the host cells we used concentrations that included the 1x MBC to maintain intra-study comparisons and doses that approximate clinically relevant ranges. We ensured that the ranges were proportionate to each other to allow comparisons to be made. For vancomycin, in this context, we used 3, 8 and 20 µg/ml whilst osteocyte exposure ranges are predicted to be 1.1-6 µg/ml and serum trough concentrations for *S. aureus* infections are recommended to be 15-20 µg/ml (20, 29). The equivalent range used herein for rifampicin was 0.75, 2.5 and 6 µg/ml. The concentrations osteocytes are expected to be exposed to are 1.3-6.5 µg/ml (20).

### Quantification of autophagic flux

Protein was isolated from differentiated cultures using M-PER^TM^ mammalian cell protein lysis reagent and Halt® protease and phosphatase inhibitor cocktail (both from Thermo Scientific, Victoria, Australia). Protein samples were quantified using the BCA protein assay method (Bio-Rad, Victoria, Australia), and 10 μg (unless specified otherwise) electrophoresed using Novex® 4–12 % gradient Bis-Tris denaturing gels (Life Technologies, Victoria, Australia) and electroblotted to Trans-Blot® Turbo nitrocellulose membranes (BioRad). Membranes were blocked in 5% diploma skim milk for 1 hour and then probed overnight at 4°C with primary antibodies directed to LC3A/B I-II (# 4108 in 5% BSA), Sequestosome (#5114 in 5% skim milk; both Cell Signaling Technology, Boston, MA), and β-actin (#A1978 in 5% skim milk; Sigma-Aldrich Co.) followed by matching horseradish peroxidase-conjugated secondary antibodies for 1 h at room temperature (R&D Systems, MN, USA). Chemiluminescent imaging was performed using the LAS-4000 platform and histogram densitometry was performed using Multi Gauge software (V3.1 Fujifilm, Tokyo, Japan). Density scores were normalised to the β-actin control and analysis performed consistent with relevant guidelines (30, 31).

### Transmission Electron Microscopy (TEM)

Uninfected SaOS2-OY cells were prepared for TEM, as previously described (6, 27). Briefly, they were cultured in 75 cm^2^ cell culture flasks before 24 hour, 37°C exposure to fixative containing 1.25% v/v glutaraldehyde, 4% w/v sucrose and 4% w/v paraformaldehyde in PBS. After fixation the cells were demineralised to facilitate sectioning in Osteosoft^TM^ solution (Sigma-Aldrich). The fixed and demineralised cells were processed for imaging in 2% w/v osmium tetroxide before being embedded in resin and cut into 5 µm sections.

### Intracellular infection escape assay

In order to assess resuscitation of intracellular *S. aureus* infections, a bacterial outgrowth or escape assay was generated. SaOS2-OY in 48-well plates were infected with a dilute suspension of WCH-SK2 (0.05 MOI; in PBS). The infection was maintained for 2 weeks in a solely intracellular state under the pressure of lysostaphin supplemented differentiation medium, as reported (27), to shift the bacterial community into a chronic-like, predominantly non-culturable state. At 15 days post-infection media were replaced with antimicrobial-free differentiation media containing the modulators of autophagy Ku-0063794 (autophagy promoter; Sigma-Aldrich), bafilomycin (autophagy blocker; Jomar, VIC, Australia) or untreated control media. Additionally at this time, host cell lysates were assessed for culturability and fewer than 8 CFU/well was measured. Following the initiation of treatment, bacterial content was assessed daily (excluding 6 days post-treatment) by drying 2 µl/well of supernatant onto NBA plates to ascertain whether bacteria with a culturable phenotype had escaped into the media.

### Statistical analyses

All statistical analyses were performed using GraphPad Prism software v. 9.0.0 (GraphPad Prism, La Jolla, CA, USA). Differences in CFU counts over time were tested using one way ANOVA with Tukey’s post-hoc test. Antibiotic dose responses analysed by Western blot were analysed by simple linear regression. Protein levels in response to various treatments were tested by Brown-Forsythe ANOVA test with unpaired t with Welch’s correction multiple comparisons test. Antibiotic dose effects on cell viability analysed by LDH assay were analysed by simple linear or non-linear regression and plotted with 95% confidence intervals. Outgrowth well survival curves were compared with the Log-rank Mantel-Cox test. In all cases, a value for *p* < 0.05 was considered significant.

## RESULTS

### Effects of rifampicin and vancomycin on *S. aureus* intracellular culturability and bacterial burden

*S. aureus* intracellular infections, using the MRSA strain WCH-SK2, were established in SaOS2-OY cells. Consistent with our published observations with this model (27), examination by TEM revealed both viable and dividing intracellular *S. aureus*, as well as bacteria in the process of being cleared by autophagolysosomal degradation (**Fig. 1A**). Cultures were exposed to rifampicin and/or vancomycin at 1x the respective MBC. In the absence of antibiotics, the SaOS2-OY model demonstrated the capacity to clear a significant proportion of intracellular bacteria over the course of the observation interval with a time-dependent decrease in recovered intracellular CFU and total bacterial abundance (**Fig. 1B**). Rifampicin treatment was effective in clearing culturable intracellular bacteria and this effect was heightened with co-exposure to vancomycin. Vancomycin treatment alone failed to cause a significant decrease in CFU/well. Despite effects on CFU, neither antibiotic alone or in combination affected intracellular bacterial DNA levels compared to untreated controls (**Fig. 1C**), suggesting bacterial persistence in the face of these treatments. Interestingly, bacteria recovered from rifampicin-treated cells displayed rifampicin resistance while those that grew from vancomycin-treated cells retained vancomycin sensitivity (data not shown).

**Figure 1.**
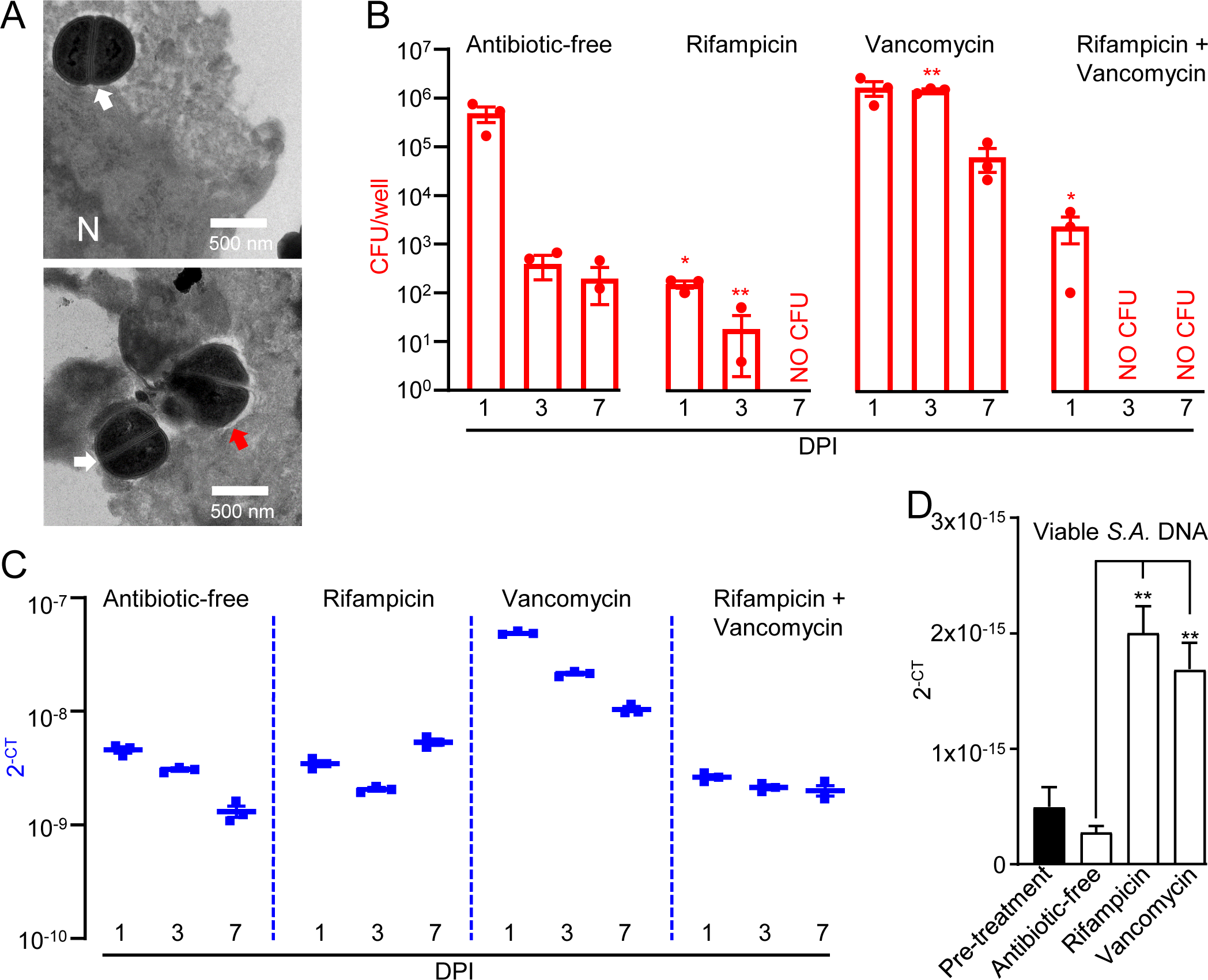
Effects of rifampicin and vancomycin on intracellular CFU and bacterial DNA levels. SaOS2-OY cultures were infected with WCH-SK2 at an MOI of 1. (A) TEM analysis at 3h post-infection (N = nucleus; white arrow: dividing *S. aureus*; red arrow: degrading *S. aureus*). Cultures were either untreated or treated with 1 x MBC of rifampicin (0.75 µg/ml), 1 x MBC of vancomycin (8 µg/ml) or a combination of both antibiotics at their respective MBC: (B) CFU counts from lysates of triplicate wells at 1, 3 and 7 days DPI in the presence or absence of antibiotic treatments were measured. Differences between treatments and antibiotic-free controls were determined at each time point using Student’s t-tests; (C) Half of the lysate from each well was pooled within groups and for total DNA isolation following treatment with the PMAxx™ reagent. Bacterial DNA abundance was measured from this by qPCR for *sigB*, expressed as 2^-CT^. (D) Effects of antibiotics in a chronic infection model were also tested in cultures infected as above but maintained in lysostaphin-supplemented media until host cell lysates yielded no CFU (17 days post-infection). Cultures were then exposed to antibiotic-supplemented (1x MBC of rifampicin or vancomycin, as above) or control media for 24 hours. Host cell lysates were then assayed for *sigB* DNA by qPCR, as above. Significant difference to controls is indicated by * *p* < 0.05 and ** *p* < 0.01. DPI = days post-infection, CFU = Colony forming units, MBC = Minimum bactericidal concentration, SA = *S. aureus*.

The potential of the antibiotics to influence the frequency of persistent intracellular *S. aureus* was further assessed in a long-term ‘chronic’ infection model. For this, SaOS2-OY cells were infected with *S. aureus* WCH-SK2 and then cultured in the continued presence of lysostaphin until measures of intracellular (and extracellular) CFU were below detection, which occurred at 17 days post-infection. Following this, antibiotic treatments were applied for 24 hours. In order to exclude molecular detection of dead bacteria, the photoreactive viability PCR reagent, PMAxx^TM^, was used. **Figure 1D** shows that non-culturable bacterial abundance was appreciably increased within the SaOS2-OY cells challenged with either antibiotic compared with controls. This outcome points to the antibiotics deleteriously interacting with the host cell innate antimicrobial defences, promoting the intracellular persistence and expansion of *S. aureus*.

### Effects of rifampicin and vancomycin on autophagic flux in osteocyte-like cells

To examine possible off-target effects of antibiotics on bacterial clearance mechanisms, SaOS2-OY cells were exposed to clinically-relevant dose ranges of rifampicin or vancomycin and assessed for alterations of autophagic flux. The SaOS2-OY model demonstrated significant autophagic potential, as indicated by the differential LC3A/B-II signal for the control exposure (i.e. basal LC3A/B-II) and its accumulation with the autophagy inhibitor, bafilomycin (+2.76-fold/flux of LC3A/B-II for bafilomycin vs no-exposure, β-actin normalised, *p* = 0.025, n = 3, **Fig. 2A-B**). The abundance of LC3A/B-II positively correlated with increasing concentrations of rifampicin and vancomycin (**Fig. 2B**, *p* = 0.044 and *p* = 0.047, respectively), and appeared to plateau at a similar proportion of maximal flux inhibition (indicated by levels in the co-presence of bafilomycin), 34% and 37%, respectively. While apparent accumulation of p62/SQSTM1 was also observed (**Fig. 2A**), the repeat outcomes did not achieve statistical significance, likely due to an already relatively high basal abundance, which was not altered even in the added presence of bafilomycin (**Fig. 2C)**. While rifampicin displayed cytotoxicity at doses above 25 µg/ml, neither antibiotic impaired cell viability in the dose-ranges utilised for protein-based or cell culture experiments (**Fig. 2D**).

**Figure 2.**
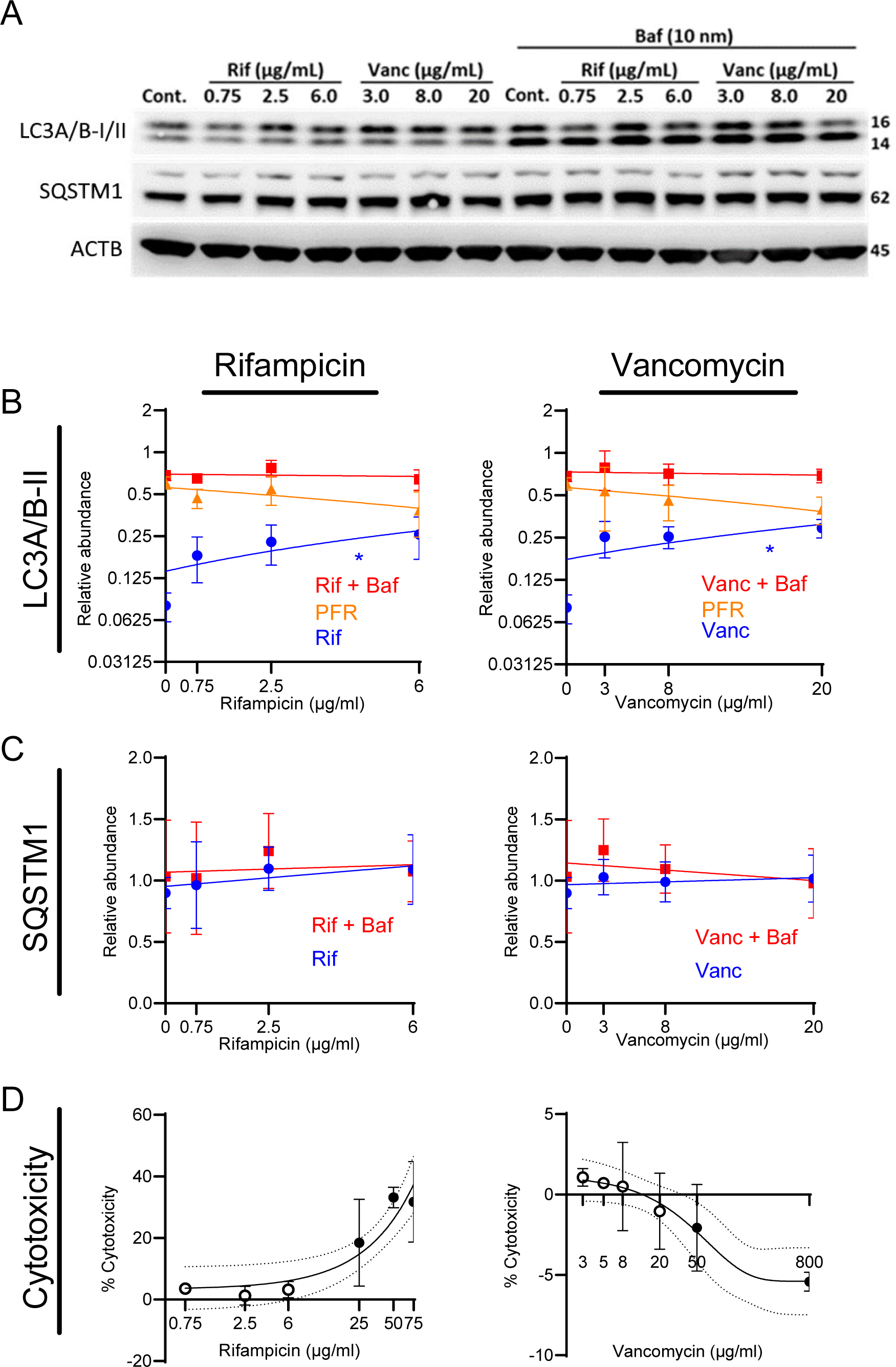
Rifampicin and vancomycin modulate autophagy in the SaOS-2 model. Relative LC3A/B-II quantity per well was tracked across respective dose ranges of rifampicin and vancomycin representative of what osteocytes *in vivo* would be expected to be exposed to clinically for either antibiotic. Measurements were taken at 6 hours following treatment initiation, which were performed either with or without bafilomycin. (A) Representative example of a blot from which data were quantified. Quantification of the relative abundance of (B) LC3A/B-II and (C) *p62/SQSTM1* with or without bafilomycin. Significance values associated with the linear regressions indicate whether the slope of the relationship was significantly non-zero. Potential flux remaining was determined by subtracting the treatment values from the treatment + bafilomycin values. (D) Percent cytotoxicity was determined using an LDH enzyme activity assay following 24 hours of exposure to a dose range of either antibiotic. Hollow data points represent doses within the range that osteocytes would be expected to be exposed to during clinical treatment *in vivo* and the dose range used within this figure. The 95% confidence interval is presented with the line of best fit, which for rifampicin was a linear regression and for vancomycin a one-phase association curve. Cont. = Control, Rif = Rifampicin, Vanc = Vancomycin, Baf = Bafilomycin, PFR= Potential Flux Remaining. Significant association between the variables are indicated by * *p* < 0.05.

### Effect of autophagic flux modulation in an acute infection model

Given confirmation of antibiotic inhibition of autophagy, we next examined whether the autophagy apparatus and *S aureus* interact within the context of an acute infection model. When SaOS2-OY cultures acutely infected with *S. aureus* WCH-SK2 were subjected to exogenous potentiation of autophagic flux with KU-0063794 (inhibitor of mTORC1 and mTORC2), there was a significant 0.12 ± 0.03-fold reduction relative to the control in the recoverable intracellular CFU/well (**Fig. 3A**, *p* = 0.015). In line with this observation, blockade of autophagy with bafilomycin resulted in a 2.1 ± 0.3-fold increase in the number of CFU within the host cell lysates (**Fig. 3A**, *p* = 0.004). However, neither autophagy modulator altered the bacterial genomic *sigB* signal (**Fig. 3B**). This indicates that a significant proportion of the bacteria may be protected from, or otherwise do not interact with, the autophagic apparatus in the host cell, or in response to exogenous modulation of autophagy, the bacteria modulate their growth phenotype.

**Figure 3.**
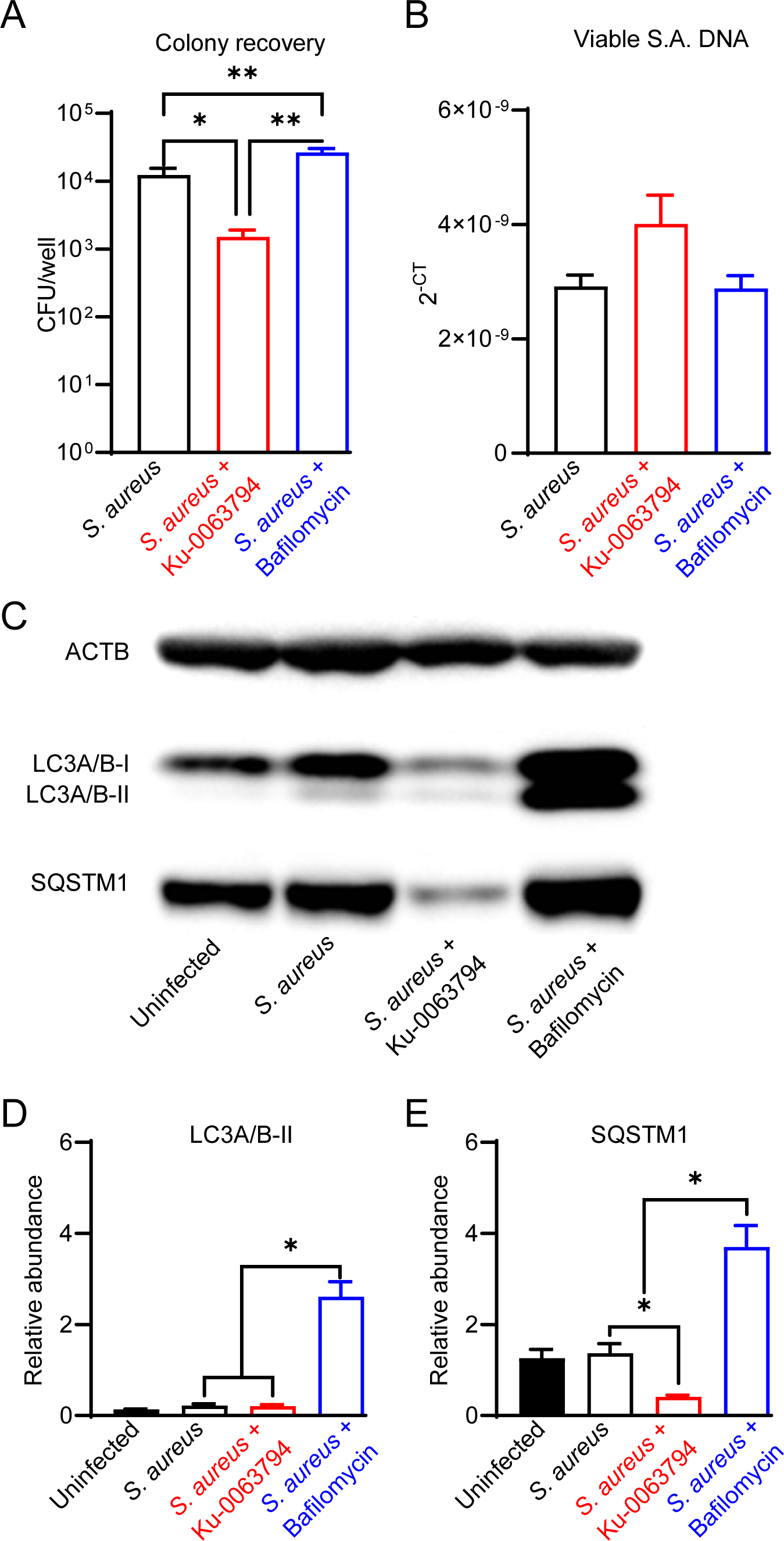
Analysis of the impact of autophagic flux promoters or blockers on bacteriological and autophagic outcomes in an acute infection context. Differentiated SaOS2-OY cells were pre-treated with either control media (no treatment) or with media containing either Ku-0063794 (4.3 µM) or bafilomycin (10 nM) for 4 h. then exposed to WCH-SK2 at MOI = 1, as described in *Materials and Methods*. After the 2 h infection period, treatments were replenished in media also containing lysostaphin to clear extracellular bacteria and cells cultured for a further 24 h before assaying for: (A) Colony forming units (CFU) per well from host cell lysates; (B) relative *sigB* DNA expressed as 2^-CT^ from host cell lysates treated with PMAxx™ reagent prior to DNA purification. (C) Representative image of a Western blot (n = 3) used for protein quantification; (D) Relative LC3A/B-II protein quantity per well from host cell lysates. (E) Relative p62/SQSTM1 quantity per well from host cell lysates. Protein levels were normalised to ACTB. Significant difference to controls is indicated by \**p* < 0.05 and \*\**p* < 0.01.

Western blot analysis of protein lysates from these cultures confirmed modulation of autophagy in an infection context (**Fig. 3C-E**). Infected controls exhibited slight, albeit non-significant, blockage of autophagy based on visually elevated LC3-II and p62/SQSTSM1. Strong induction of autophagic flux in infected cells was seen with Ku-0063794, which resulted in significantly reduced p62/SQSTSM1 (*p* < 0.05). Flux potentiation also resulted in apparent reduced LC3A/B-II abundance relative to the infected control but this did not achieve significance, likely due to the already low starting level. Bafilomycin treatment resulted in the concomitant accumulation of these markers of autophagic flux, consistent with its inhibition.

Together, these results suggest that while a proportion of the infecting bacteria is susceptible to autophagic attack, which can be increased by exogenous potentiation of autophagy, a consistent and substantial sub-population of viable *S. aureus* remain within the host cell.

### Effect of autophagic flux modulation in a chronic infection model

We next investigated the influence of autophagy modulation in a chronic infection model, where the infected cultures first exhibited intracellular bacteria with a uniformly non-culturable phenotype prior to treatment, i.e. at 8-days post-infection. Promotion of autophagy in chronically infected cultures with Ku-0063794 maintained the culturable and viable population to a similarly low level as the control exposure (*p* = 0.26), as expected. However, inhibition of autophagy with bafilomycin resulted in a small but significant increase in culturable intracellular bacteria, consistent with autophagy actively clearing *S aureus* during the observation interval (**Fig. 4A**). Similar to the acute model, neither treatment affected *S. aureus* intracellular DNA levels (**Fig. 4B**). Indeed, based on relative DNA levels, approximately the same frequency of an apparent persistent sub-population of *S. aureus* remained between these models, despite the increased duration of infection (**Fig. 3B** vs **Fig. 4B**). Infected controls displayed blockage of autophagic flux with significantly elevated LC3A/B-II (**Fig. 4C-D**). Potentiation of autophagy resulted in a clear reduction in p62/SQSTM1, while blockage with bafilomycin increased both LC3A/B-II and p62/SQSTM1 levels (**Fig. 4C-E**). Together, these data suggest a continued and increased disassociation between autophagic flux modulation and *S. aureus* persistence in a chronic infection.

**Figure 4.**
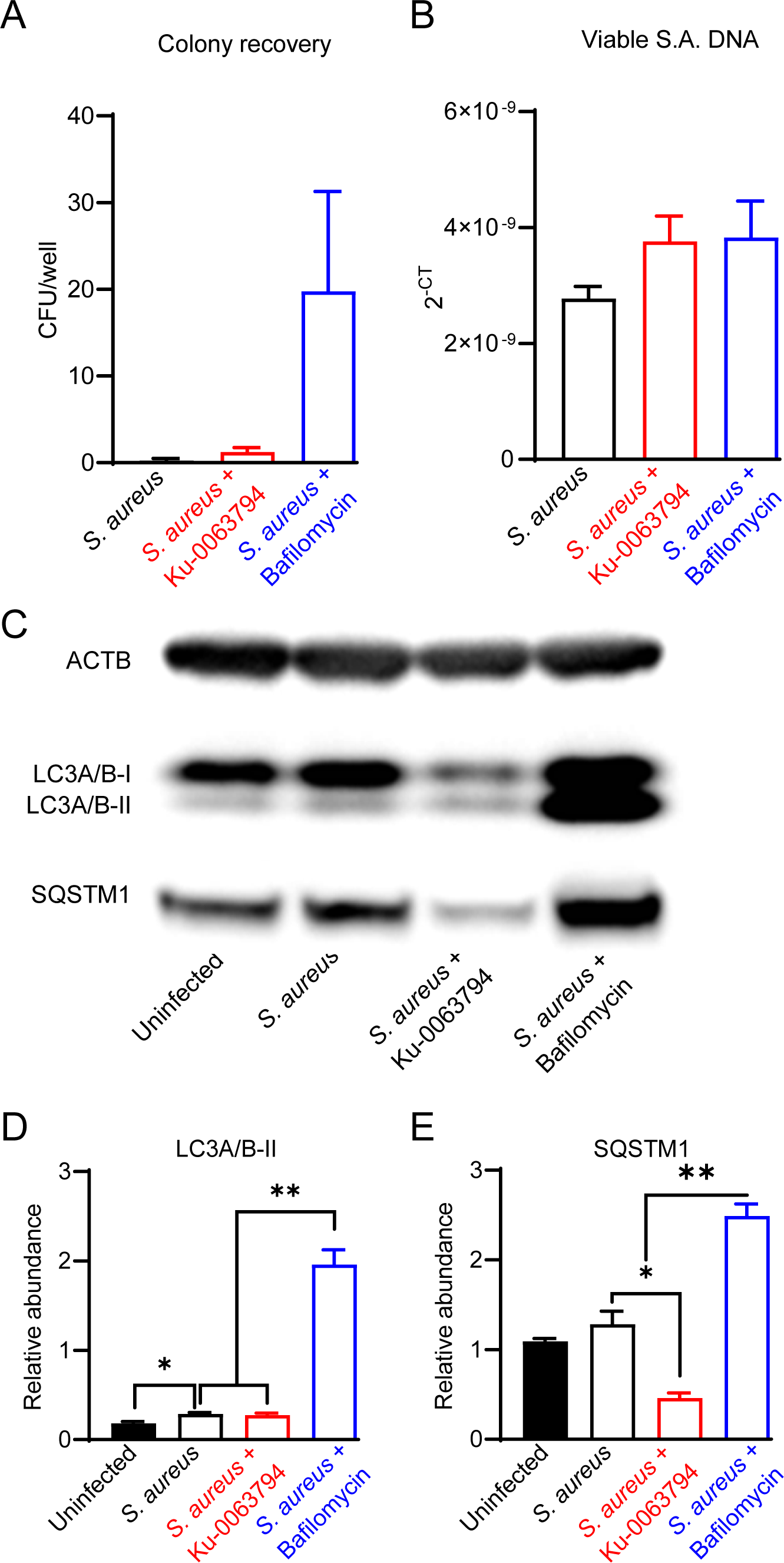
Analysis of the impact of autophagy modulators on bacteriological and autophagic outcomes in a chronic intracellular infection context. Differentiated SaOS2-OY cells were infected at MOI of 1 with WCH-SK2, as described in *Materials and Methods*. Cells were cultured for 7 days and cell lysates confirmed to yield no CFU after 24 hours. Cultures (at day 8) were then treated with either control media (no treatment) or with media containing either Ku-0063794 (4.3 µM) or bafilomycin (10 nM) for a further 24 h before assaying for: (A) Colony forming units (CFU) per well from host cell lysates; (B) relative *sigB* DNA expressed as 2^-CT^ from host cell lysates treated with PMAxx™ reagent prior to DNA purification. (C) Representative image of a Western blot (n = 3) used for protein quantification; (D) Relative LC3A/B-II protein quantity per well from host cell lysates. (E) Relative p62/SQSTM1 quantity per well from host cell lysates. Protein levels were normalised to ACTB. Significant difference to controls is indicated by \**p* < 0.05 and \*\**p* < 0.01.

### Effect of autophagic flux modulation on bacterial release

To further test the involvement of autophagic flux in bacterial persistence in osteocytes, we designed a bacterial outgrowth assay, where a long-term (17 day) non-culturable infection was treated with autophagy modulators in antimicrobial-free media. The outcome measured was detectable bacterial growth in culture supernatants, analogous to relapse of a chronic infection. This was scored as the percentage of wells containing bacterial outgrowth over time. Autophagic potentiation with Ku-0063794, when compared to blockade with bafilomycin, increased the rate of bacterial escape and growth over time, with the divergent effect becoming evident after approximately 4 days post-treatment (**Fig. 5A**, *p* = 0.03). Media from all wells with outgrowth generated confluent growth on agar plates. Where single colonies were discernible, no differences in colony morphology was seen (**Fig. 5B**): all groups presented with unpigmented colonies after 24 hours of plate incubation, which reverted to gold pigmentation and haemolytic capability upon further incubation, typical of *S. aureus* wild-type growth characteristics. This suggests that in a long-term chronic infection context, potentiation of autophagy can increase the rate of switching from a quiescent to an active growth phenotype.

**Figure 5.**
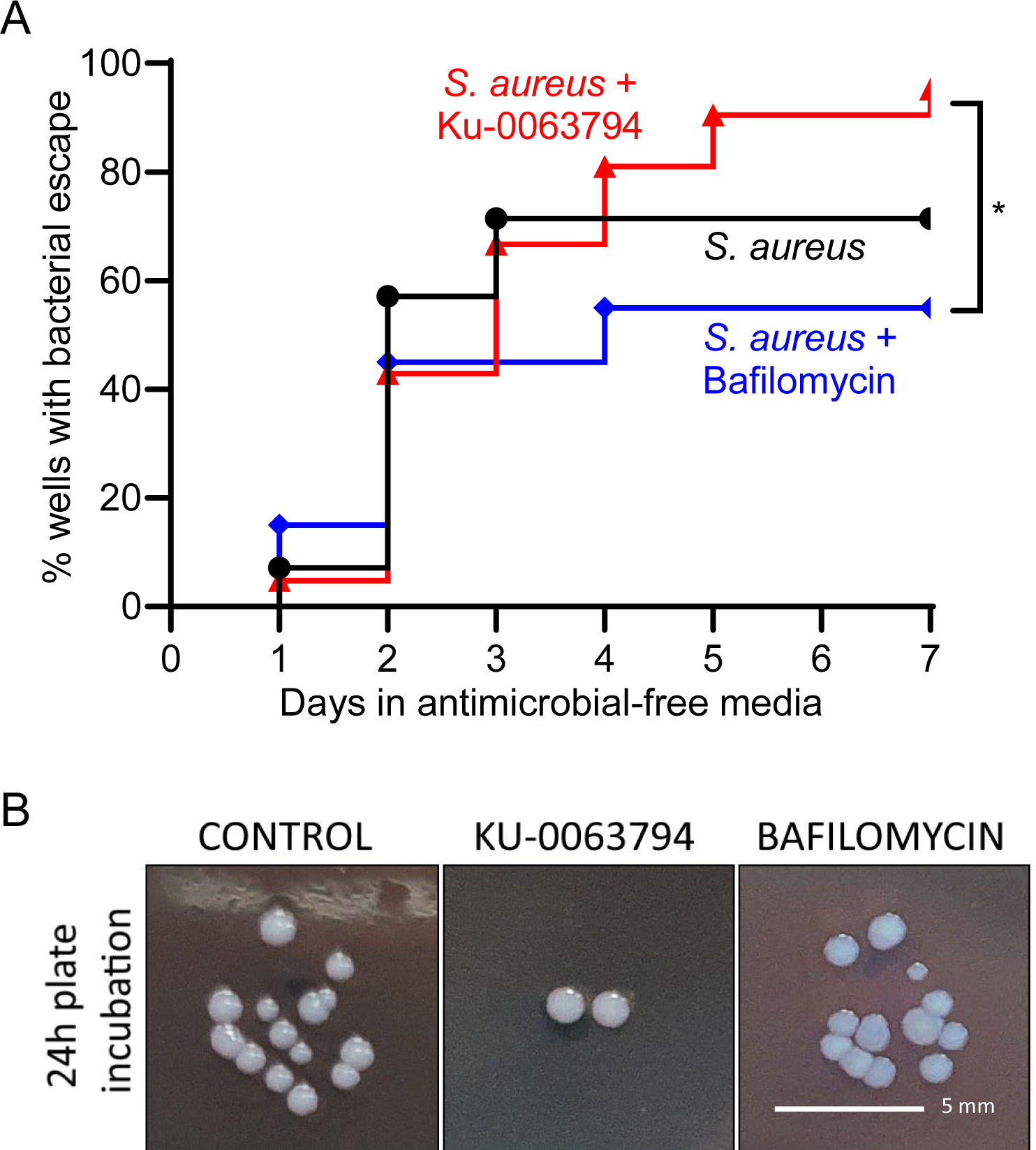
Effect of autophagic modulation on bacterial escape from the intracellular niche. Differentiated SaOS2-OY cells were infected at MOI of 1 with WCH-SK2, as described in Materials and Methods. Cells were cultured in medium containing lysostaphin for 16 days and cell lysates confirmed to yield no CFU after 24 hours. Medium were then replaced with either control medium (no treatment) or with media containing either Ku-0063794 (4.3 µM) or bafilomycin (10 nM), all in the absence of any other antimicrobial treatment, and cultures monitored by visual inspection for bacterial outgrowth for a further 7 days. (A) Inverted Kaplan-Meier estimator tracking the percent of wells with outgrowth. (B) Gross colony morphological assessment from each experimental group after 1 day of plate incubation. Statistical differences between survival curves were analysed with a log-rank test and results are indicated by \**p* < 0.05.

## DISCUSSION

Consistent with our previous reports, human osteocyte-like SaOS2-OY cells displayed the capacity to clear a significant proportion of internalised *S. aureus* in the absence of exogenous intervention (6, 27), suggesting an active and effective autophagy/xenophagy system in these cells. As the infection of osteocytes by *S. aureus* is proposed to represent an important niche of persistence within bone in PJI (6, 32), we sought to test to what extent, if any, this process was dependent on manipulation of the autophagy/xenophagy pathway. At the same time, it was important to establish if clinically utilised antibiotics were effective in this model and if off-target effects on autophagic flux influenced the treatment. Whilst rifampicin alone and in combination with vancomycin was effective in reducing recoverable intracellular CFU, we found that neither antibiotic cleared intracellular bacteria as measured by viable DNA levels, and in fact increased this measure in a chronic infection model where there was a complete absence of culturable bacteria. Neither vancomycin or rifampicin were able to eliminate viable intracellular *S. aureus* at the respective MBCs for the strain tested under planktonic conditions, which fall within the ranges we expect osteocytes to be exposed to *in vivo* during clinical use (20). This was not altogether surprising, given the additional protection the intracellular environment provides the invading pathogen. Vancomycin targets cell wall synthesis and is generally effective against *S. aureus* infections. However, its intracellular effectivity has generally been found to be poor, due in part to its slow accumulation and possibly also the natural resistance of slow growth phenotypes to this killing modality (20). Rifampicin, on the other hand, has excellent intracellular penetrance and inhibits bacterial transcription, although its effectivity against intracellular *S. aureus* appears to be strain-dependent (20). It is interesting that, whilst there were differences in the efficacy of the two antibiotics at reducing culturability, both were capable of causing an increase in the total bacterial population size in a manner that would be overlooked by current routine clinical diagnostic procedures, even though they operate through entirely distinct mechanisms.

We showed recently that enumeration of *S. aureus* by culture alone in an intracellular context lacks accuracy, certainly in human osteocytes (33) but likely also in other host cell types. *S. aureus* has been shown to replicate intracellularly in the endosomes of various cell types, including THP-1 macrophages and in HUVEC (34). Consequently, our observations suggest either that antibiotic-driven bacterial adaptation facilitates greater rates of replication within the intracellular niche, or alternatively, an effect of the antibiotics to inhibit host cell clearance of bacteria or inhibit their replication.

The presence of a non-culturable but metabolically active persister phenotype of *S. aureus* was evidenced by the relative increase in bacterial DNA within intact bacteria following treatment with both rifampicin and vancomycin, as well as the ability of a chronically non-culturable infection to spontaneously revert to a culturable state when antimicrobial pressure was removed (see **Figs. 2 & 5**). Whilst the exact nature of these persister-like cells was not characterised herein, this reflects the viable but non-culturable state often described in the context of biofilm formation (35, 36).

The effects of either antibiotic could not be explained by cytotoxicity at the doses tested. However, both antibiotics caused an acute, significant dose-dependent inhibition of autophagic flux, at their respective clinically relevant doses. Given the observed antibiotic-mediated increase in LC3A/B-II, the underlying mechanism is likely at least partial inhibition or disruption of lysosomal maturation or autophagolysosomal degradation or fusion, similar to the effects of the definitive autophagy blocker, bafilomycin (37). While the apparent lack of modification of p62/SQSTM1 quantity by bafilomycin treatment in uninfected SaOS2-OY cells implies that this adaptor protein is not involved in the basal flux being investigated, *SQSTM1* is known to be highly transcriptionally regulated (38) and a subsequent effect on protein levels may have been missed due to the time intervals examined.

Lysosomal maturation and autophagosome-lysosome fusion represent key steps in the xenophagic pathway, which facilitates the clearance of intracellular bacteria, and *S. aureus* has previously been shown to be capable of inducing the formation of a replicative niche through the dispruption of these processes in other cell contexts (39, 40). *S. aureus* indeed induced some degree of autophagic flux inhibition in our osteocyte infection model, best evidenced by the increased level of LC3A/B-II in the chronic model (see **Fig. 4**). Interestingly, p62/SQSTM1 was not significantly increased by infection itself. Potential explanations for this observation are functional redundancy via other autophagy adaptors, a proportional increase in the rate of p62/SQSTM1 degradation, additional effects on transcriptional regulation of *SQSTM1* (38) masking changes in protein levels, or via a yet to be identified effect exerted by the infecting bacteria. Nonetheless, p62/SQSTM1 appeared to retain its function, as the autophagy potentiators and blockers were capable of significantly altering relative p62/SQSTM1 quantities when bacteria were present (see **Figs. 3** & **4)**. We sought to investigate whether the paradoxical decrease in culturability but retention or increase in bacterial DNA levels observed with antibiotic treatment was explainable by autophagy dysregulation. As such, we tested the effects of bafilomycin and the autophagic flux potentiator Ku-0063794 on microbiological outcomes in intracellularly infected cells. Neither treatment when applied either immediately following infection or after chronicity was established resulted in a significant change in the quantity of viable bacterial DNA, whilst culturability was significantly modulated, suggesting effects on bacterial growth phenotype beyond effects on xenophagic clearance.

In the period immediately following infection, bafilomycin treatment caused a 2.1-fold increase in CFU recovered from cell lysates, presumably by reducing xenophagic stress on bacteria. This also occurred in a chronic infection model, exhibiting mostly non-culturable intracellular bacteria, where bafilomycin treatment appeared to facilitate a minor bacterial sub-population returning to a culturable state. However, our results when modelling recurrence of a longer-term chronic infection showed that the potentiation (but not blockage) of autophagic flux was associated with a greater rate of bacterial escape and growth into the culture media, an effect that became evident 4-7 days after treatment. These findings suggest that in the chronic, non-culturable state, potentiation of autophagic/xenophagic flux imparts selective stress for a persister, high-growth phenotype. Our findings are also consistent with the report that stimulation of autophagy promoted *S. aureus* replication and host cell killing (22). Conversely, blockage of flux may select for low-growth phenotypes. Notably, there were no gross differences in colony morphology between conditions in the outgrowth assays, suggesting that the revertant clones were not SCVs but rather persister-type cells. The possible explanation for the interaction of autophagy with *S. aureus* in the context of flux blockers or potentiators is summarised in **Figure 6**. Of note, xenophagy acting as a stress that induces a change in the growth phenotype is conceptually similar to reports of *S. aureus* adapting to autophagy/xenophagy in other host cell contexts (22, 41). This has interesting implications for the mechanisms of recurrence in dormant infections, whereby dysfunction of autophagy in the host cell contributes to the regeneration of a culturable subpopulation.

**Figure 6.**
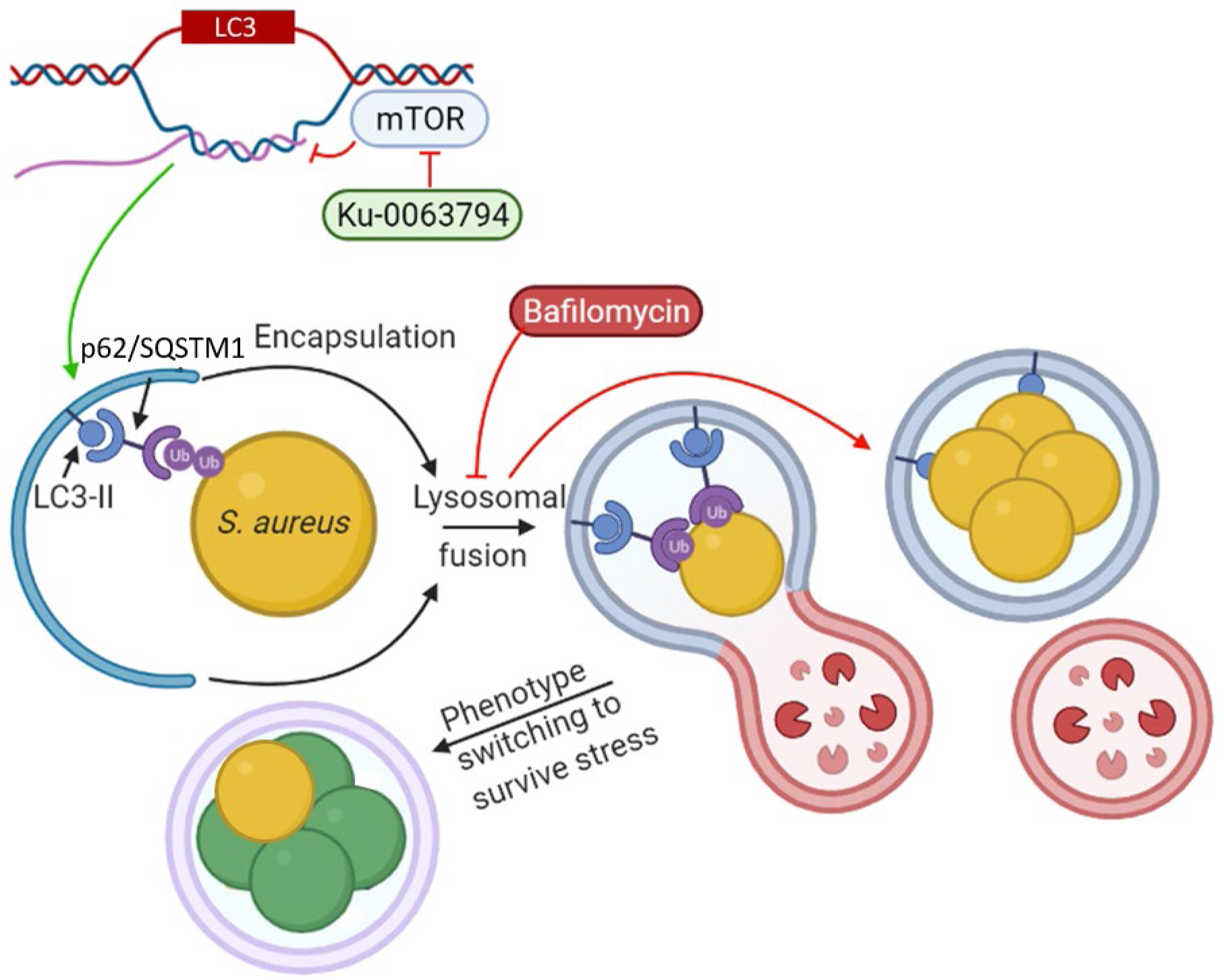
Schematic of the proposed interaction of Ku-0063794 and bafilomycin-mediated modulation of autophagy on intracellular *S. aureus* within SaOS2-OY cells. *LC3* is given as an example of an autophagic gene whose transcription is modulated by mTOR blockade. Following phagosomal entrapment and LC3A/B-II-mediated formation of the phagolysosome, *S. aureus* phenotype switching occurs (*green* cells) resulting in a decline in culturability on nutrient agar, although with a sub-population of persister cells (*gold*) with potential to revert to a wild-type growth state, such that there is no significant change in bacterial population size. This process is promoted through xenophagic potentiation with Ku-0063794. Treatment with bafilomycin prevents lysosomal fusion promoting a culturable phenotype. Ub = Ubiquitin. *Image created using Biorender*.

Limitations of this study include that xenophagic activity was measured by proxy via the flux of LC3A/B and p62/SQSTM1; this targeted analysis would miss potential changes in other pathways and proteins involved in xenophagy. Furthermore, in future studies, temporal and/or unbiased transcriptional regulation of autophagy markers also should be taken into account. Bacterial outgrowth was recorded as percentage of culture wells that became positive for growth; it was not possible to determine how many revertant events took place in any single well and this could have varied between treatments. In addition, the scope and clinical applicability of the observations is limited by the fact that a single *S. aureus* strain was used within a cell line model. This was done for consistency between readouts and it is possible that other clinical *S. aureus* isolates might respond differently. The use of SaOS2-OY was also used for consistency, as donor variability is a potential confounder with primary cell models, however this is the only validated human osteocyte-like cell line model described to date.

In conclusion, rifampicin alone and vancomycin treatment of human osteocyte-like cells caused some degree of autophagic flux inhibition, as did infection of these cells with *S. aureus*. Rifampicin and vancomycin affected the proportion of culturable bacteria but either did not affect the total population size or increased it. Targeted autophagic flux modulation in both the acute and chronic-like infection contexts tested were also capable of significantly affecting the proportion of culturable intracellular *S. aureus* but not the size of the intracellular bacterial pool. This suggests that a persister population of *S. aureus* escapes both antibiotic and autophagic/xenophagic stress by adopting a non-culturable growth phenotype. In contrast, in a long-term infection, potentiation of autophagy promotes *S. aureus* reactivation and escape from the host cell, whereas inhibition of autophagy appear to promote an intracellularly resident growth phenotype, suggesting that certain disruptions to host cell metabolism could be stimuli for recurrence of intracellular *S. aureus* infections.

## ACKNOWLEDGEMENTS

NJG and AZ were supported by an Australian Postgraduate Award and Divisional Postgraduate Scholarship from the University of Adelaide, respectively. ER received support from The Royal Adelaide Research Committee, Royal Adelaide Hospital Research Fund, The Health Services Charitable Gifts Board, and the Rebecca L Cooper Medical Research Foundation. This work was supported by funding to GJA and DY from the National Health and Medical Research Council of Australia (NHMRC, Grant ID 2011042).

