## Supplemental Table 1 for "*Staphylococcus aureus* Persistence in Osteocytes: Weathering the Storm of Antibiotics and Autophagy/Xenophagy"

Supplementary Table 1. PCR primers to amplify genomic DNA sequences.

| Gene | Primer direction | Oligonucleotide Sequence (5'→ 3') |
|---|---|---|
| ACTB | forward | CGCGAGAAGATGACCCAGATC |
| | reverse | TCACCGGAGTCCATCACG |
| sigB | forward | GGGGCAACAAGATGACCATT |
| | reverse | TGCCGTTCTCTGAAGTCGTG |
